# Quantifying the predictability of evolution by analysis of coalescent rate variation

**DOI:** 10.1101/2025.09.29.679185

**Authors:** Erik Volz, Xavier Didelot

## Abstract

We investigate the predictability of evolution in terms of the phylogenetic placement of new lineages. This leads us to develop a class of coalescent models that relax neutrality by allowing the rate of coalescence to vary as a continuous heritable trait. In this setting, each lineage has a relative propensity to coalesce, with coalescent odds ratios defined from the product of pairwise propensities. Estimated coalescent odds provide a statistic that captures variation in lineage growth and are informative about the strength of natural selection acting on individual lineages. A number of practical statistical methods are then developed: techniques to adjust for biased and non-uniform sampling; procedures to automatically calibrate hyperparameters governing the evolution of coalescent propensity; and, methods for clustering phylogenies into sets that delineate clades according to coalescent propensity. Simulations show sensitivity of these methods to detecting small selective effects acting on rare variants, and strong robustness to imbalanced sampling. We demonstrate these methods using two datasets reflecting microbial populations evolving under strong selection. First, we examine a large set of *Neisseria gonorrhoeae* genomes, and show that lineages with high coalescent odds feature a unique antibiotic resistance pattern which presaged its subsequent expansion. We then re-analyse SARS-CoV-2 data in combination with independent estimates of reproduction numbers corresponding to major variants of concern 2020-2023. This indicates that coalescent odds can function as an excellent tree-based proxy for relative fitness of major SARS-CoV-2 lineages.

## 1 Introduction

Molecular evolution can be predicted in situations where distinct genetic variants co-circulate and are governed by selective forces that can be estimated [1]. In the context of phylogenetics, the predictability of evolution may be framed in terms of the capacity to predict the phylogenetic placement of taxa collected in the future. For example, every branch in a phylogenetic tree reconstructed from molecular data corresponds to a set of shared genetic features (mutations, indels, etc.) of the clade descended from that branch. If these features are not neutral, future samples are more likely to be descended from the branch representing the evolution of fitness-enhancing features. A phylogenetic approach to quantifying selective differences and predicting evolution has the advantage of encompassing all genetic diversity of the sample used to reconstruct a phylogeny, and allows the study of fitness variation both within and between clades of interest.

In this work, we examine the problem through the lens of coalescent theory. We consider models which allow coalescent rates to vary continuously across a tree as if the propensity to coalesce was a heritable trait. This has resonance with previous work on the *multiplicative coalescent* [2] which also models correlation in coalescent rates along lineages in a genealogy. But, the focus of this study is on the development of statistical methods to both detect variation in coalescent rates and to estimate its magnitude. We will not propose a generative model for trees with coalescent rate variation, but will develop a statistical model for inferring latent variables that quantify coalescent rate variation and treating the phylogeny as data. Our model for rate variation is more flexible than described in [2], encompassing situations where rates can both increase and decrease as a path is followed through a genealogy, and rates are not directly related to the number of descendants of each lineage.

In terms of the applications of the methods we develop, there are parallels with heuristic methods such as the local branching index (LBI) [3], a statistic defined for each node in a phylogeny, and defined as the exponentially weighted tree length surrounding the node. It has been widely applied in the study of virus evolution, such as for vaccine strain prediction. These applications have demonstrated some capacity to predict phylogenetic evolution purely from the topology of phylogenetic trees and without any additional information about the phenotype of sampled taxa, predictability based on this metric is usually fleeting [4, 5]. Methods such as the LBI and related phylogenetic clustering methods also depend on subjective hyperparameters, such as the rate parameter used to define exponentially-weighted tree length. They are very sensitive to imbalanced sampling, as the LBI is always higher in subpopulations that are highly sampled. We therefore aim to develop a theoretically-grounded method which is relatively robust to imbalanced sampling and to provide principled methods for calibrating hyperparameters.

To the extent that new methods can characterise fitness variation across a phylogeny, there is utility for systematics and for detecting unrecognised variation in phenotype or population structure. Phylogenies often show measurable structural deviations in terms of their topology and branch lengths from theoretical predictions based on neutral evolution. This can confound phylodynamic studies which are typically based on neutral evolution models, but can also be an object of study in its own right [6, 7]. Very often, there is inadequate data to adjust for such variation, such as if variation in fitness has not or cannot be measured independently from genetic data. Therefore, there is substantial value in methods for the detection and quantification of population structure or differences in fitness from phylogenetic data and in the absence of other metadata or phenotypic data.

## 2 Methods

### 2.1 Continuous variation of coalescent propensity

Suppose that at time *t* before present there is a number *A*(*t*) of extant lineages. Any pair of these may coalesce to form a new node in the tree. In neutral population genetic models, every lineage counted in *A*(*t*) would have equal probability of coalescing conditional on a coalescent event occurring [8]. Similarly, in a birth-death framework, every lineage would have equal probability of branching conditional on a branching event occurring [9].

Here we will relax neutrality by allowing every lineage *i* to have a relative propensity *ρ*_i_ ∈ ℜ^+^ of coalescing. If lineages *i* and *j* > *i* are extant at the time of the *k*-th coalescent event then the probability that *i* and *j* coalesce at this event is *P*_*ijk*_ ∝ *ρ*_*i*_*ρ*_*j*_. Using some approximations that are made explicit below, the variable *ρ*_*i*_ can be interpreted as the odds ratio that the branch *i* coalesces relative to co-circulating lineages. Our goal is not to propose a mechanistic process by which *ρ* varies across a genealogy as in the structured coalescent and structured birth-death models [10, 11]. Rather, we will propose a phenomenological statistical model that allows us to estimate variation in *ρ* from data.

The statistical models are premised on data of the following form: a binary tree with *n* tips and *M* = 2*n* − 1 nodes (including tips); each node has an associated ordinal index with the first *n* indices associated with tips. The last index *M* is associated with the root of the tree. Each node *i* has an associated time before present *t*_i_. If *i* ≤ *n, t*_i_ represents the known time of sampling; this time axis has a retrospective orientation, so that the time of an ancestor exceeds the time of a descendent. We further impose a time order on internal nodes, so that *t*_i+1_ > *t*_i_ for all *i* > *n*. The tree may be heterochronous, which means sampling may be at different time points. Associated with each node *i* < *M* (excluding the root) is a branch to the ancestor node *a*(*i*) which has length *l*_i_ = *t*_a(i)_ − *t*_i_. While this format of a genealogical tree is compatible with standard coalescent and birth-death models and is chosen for mathematical exposition, the statistical framework for estimating *ρ*_i_ is easily extended to time-scaled trees with multifurcations [12]. Since each node has an associated time, we can define the set of *co-circulating lineages* at time *t* to be the set A(*t*) = {*j* ∈ {1, …, *M* − 1}|*t*_*j*_ ≤ *t, t*_*a*(*j*)_ > *t*} of size |A(*t*)| = *A*(*t*). The terms “node” and “lineage” are used interchangeably.

As previously noted the probability for any two lineages *i* and *j* > *i* that are extent at the time of coalescent event *k* is *P*_*ijk*_ ∝ *ρ*_*i*_*ρ*_*j*_. We can now write the full expression for *P*_*ijk*_ by considering all possibilities of pairs of coalescing lineages and that they need to sum up to one:

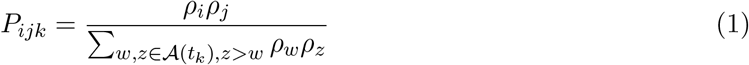

Note that if *ρ*_*i*_ = 1 ∀*i* ∈ {1, …, *M* − 1}, the neutral coalescent is obtained, since for any two extent lineages:

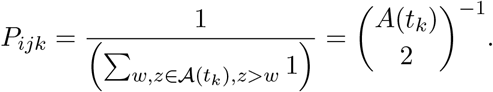

Most of the following exposition will work with the log-transformed coalescent propensities *ψ*_*i*_ = log(*ρ*_*i*_). Working with the variables *ψ*_*i*_ with support on the real line will allow us to model the evolution of coalescent probabilities across the tree as a random walk (c.f. Section 2.2).

Let us write the probability of the *i*-th lineage coalescing at the *k*-th event by marginalising over co-circulating lineages *j* and the two orders in which *i* and *j* can appear:

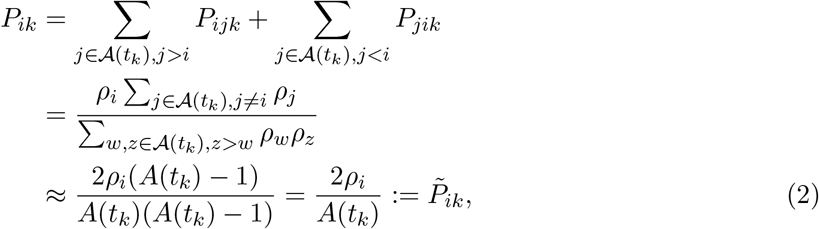

where the approximation applies if *A*(*t*_*k*_) is large and using E[*ρ*] = 1, since *ρ* represents the relative odds of coalescing. The logit transform of this probability is then

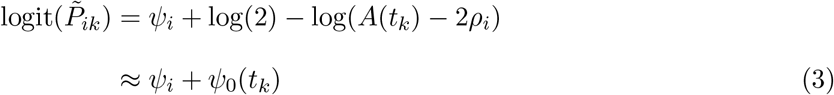

where *ψ*_0_(*t*) is a time-dependent intercept term

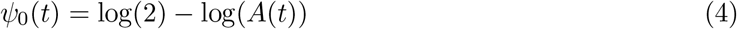

This approximation is valid when *ρ*_*i*_ ≪ *A*(*t*). Framed in this way, we can see that estimating coalescent propensities is related to fitting a logistic regression with coefficients for each lineage and a time-dependent intercept. This connection is made explicit in Section 2.3.

Before proceeding with definition of the statistical model, it is worth considering the properties of the coefficients *ψ*_*i*_ in a given tree. In a neutral model (e.g. Kingman coalescent or birth-death model), the coalescent log odds should be *ψ*_*i*_ = 0 for all *i*. Any measurable deviation from zero is evidence for population structure, biased sampling, or natural selection.

We also note that this model could be defined alternatively in terms of variation in coalescent rates, rather than variation in coalescent odds. Specifically suppose the pairwise coalescent rate is *λ*(*t*) = 1*/N*_e_(*t*), with time-varying effective population size *N*_e_(*t*). We can define the rate that branch *i* coalesces as *λ*_*i*_(*t*) = *λ*(*t*)(*A*(*t*) − 1)*ρ*_*i*_. Thus, modelling variation in coalescent odds leads naturally to an estimate of coalescent rates provided *N*_e_(*t*) can be estimated, however with our model defined in terms of the logistic-scaled *ψ*_*i*_, this is optional rather than a requirement. It is in fact a useful feature of this model that variation between lineages in coalescent odds can be estimated independently of coalescent rates which would additionally depend on population dynamics and bottlenecks [13, 14, 15].

### 2.2 Models for coalescent rate variation

With *P*_*ijk*_ defined for each coalescent event (Eq. 1), we can compute the likelihood of the entire tree. But a statistical model must further specify the correlation structure between variables. Here we propose a Gaussian Markov Random Field (GMRF) model [16] for the variables *ψ*_*i*_ which induces a tunable correlation between values on neighbouring branches. The design of this model is motivated by the necessity of capturing the empirical effects of heritability of branching rates in phylogenies without specifying an explicit mechanism that induces this correlation. For example, coalescent odds may be correlated due to natural selection (the appearance of a highly-fit variant), founder effects, or design (e.g. directed evolution). Disparities in coalescent rates can also be the consequence of biased or skewed sampling (e.g. over-sampling a geographic region), and these methods will also suggest a way to detect such analytical artefacts (c.f. Section 2.5).

According to this model, given a non-root lineage *u* < *M* with immediate ancestor *a*(*u*) and branch length *l*_*u*_, the difference in coalescent odds between *a*(*u*) and *u* is distributed as

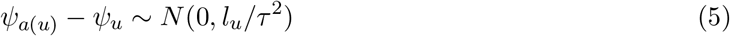

where *τ* is a *precision* parameter which must be optimised or manually set according to external criteria. We denote the likelihood of the GMRF model

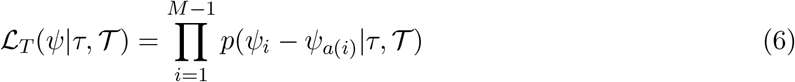

where T represents the phylogenetic data which defines branch lengths and genealogical relationships.

Let the sets *Y*_k_ = {*i, j*} contain the two nodes coalescing at the *k*-th coalescent event, that is, so that *a*(*i*) = *a*(*j*). The likelihood of the observed coalescent events can be written:

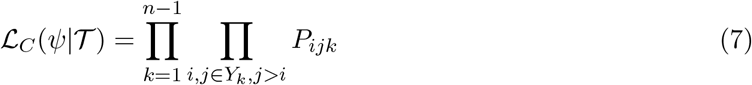

The complete likelihood of this model is

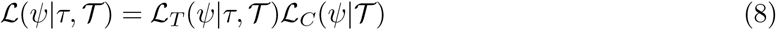

The parameters *ψ* can be estimated by maximum likelihood (ML) conditional on *τ* . A different criterion is required for selecting *τ* (c.f. Section 2.4), and the optimal value of *τ* may be application-specific. We implemented a ML procedure using the Broyden-Fletcher-Goldfarb-Shanno (BFGS) gradient descent algorithm [17] with initial conditions selected using the least squares approach in the next section.

### 2.3 Weighted least squares approximations for scalable inference

Direct optimisation of the likelihood (Eq. 8) is infeasible for many large phylogenies since the number of parameters to be estimated is 2*n* − 2. Furthermore, our methods for calibrating *τ* and adjusting for sample bias (c.f. Section 2.5) require multiple optimisations over a range of hyperparameters. Therefore it is important to find approximations to the likelihood that enable faster optimisation.

In developing fast approximations, it is helpful to define the indicator variable *y*_*ik*_ for the *k*-th coalescent event:

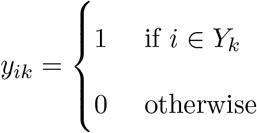

We first consider an approximation based on a scenario where the events that all lineages *i* in *A*(*t*_*k*_) coalesce are mutually independent; that is, the probability that *l* ∈ *A*(*t*_*k*_) coalesces at the *k*-th event is independent from the event that *i, j* ∈ *A*(*t*_*k*_) coalesce at the *k*-th event. This is of course a departure from the coalescent model since it unrealistically includes the possibility of other than two lineages (including zero) coalescing at the *k*-th event, however it yields a more tractable binomial likelihood, and we can quantify the difference with the true likelihood:

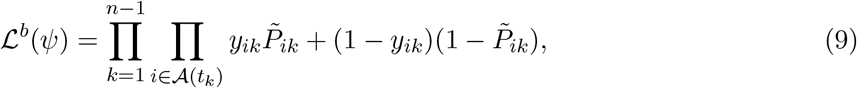

where 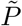 is the marginalised likelihood of each lineage coalescing from Eq. 2. Gathering terms, this becomes

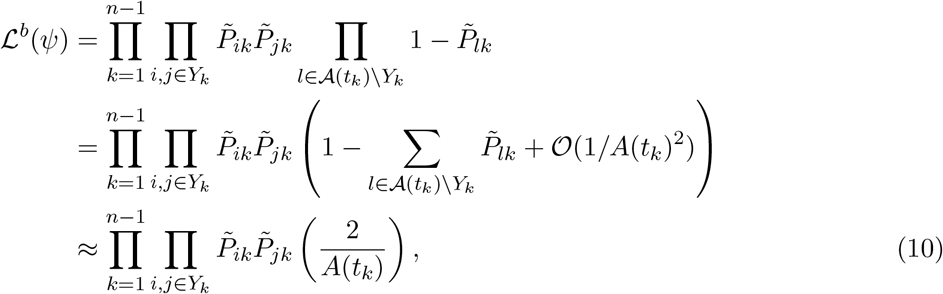

where the last approximation comes from excluding *O*(1*/A*(*t*_*k*_)^2^) terms and using 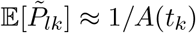, and noting that the sum has *A*(*t*_*k*_) − 2 terms. The factor Π_*k*_ 2*/A*(*t*_*k*_) is a multiplicative constant to this likelihood, so Eq. 10 will be approximately proportional to the correct likelihood (Eq. 8) providing that sample sizes are large (so that *A* is large) and where *ρ*_*i*_ ≪ *A*(*t*_*k*_) for most of the data. This approximation would fail for small trees or if there is extreme variance in coalescent odds.

Recalling that the logistic transform of 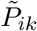 is defined in terms of the coalescent log odds *ψ*_*i*_ (Eq. 3), optimising the likelihood in Eq. 10 is equivalent to fitting a generalised additive logistic regression model (GAM) with a GMRF smooth on the estimated parameters *ψ*. In this case, the parameters *ψ* are treated as coefficients on observations corresponding to lineages that are extant at each event. This furthermore requires pre-specification of the precision parameter *τ* (see next section) and must make use of the approximate intercept expressed in Eq. 4. Such a regression model can be readily fitted using e.g. the `mgcv` R package [18].

Fitting the binomial model (Eq. 10) as a GMRF-GAM is an improvement in speed over direct optimisation of Eq. 8, although this will still be computationally expensive for large trees. We therefore consider another approximation based on linearising the binomial likelihood around *ψ* = 0, which leads to a weighted least squares estimator. Consider the log likelihood of a single term of the likelihood in Eq. 9 corresponding to node *k*, and making the dependence of 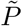 on *ψ* clear:

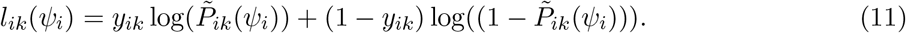

Using our definitions in Eq. 4, so that derivatives are: 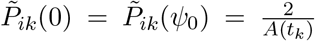, we have that the first two derivatives are:

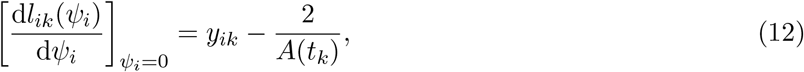

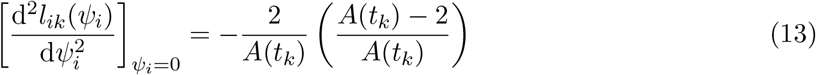

Using a 1-step iteratively reweighted least squares (IRLS) method [19], we can estimate *ψ* by regression on the pseudo-data:

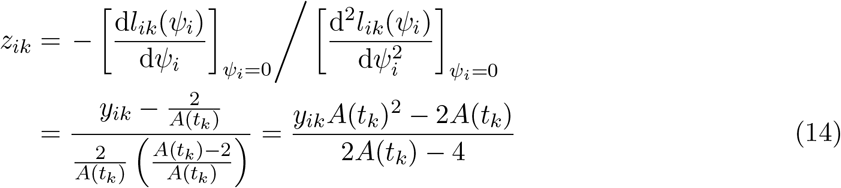

The weight corresponding to each term comes from the binomial variance:

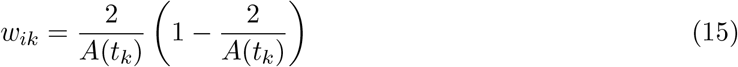

In order for the least squares solution to approximate the full likelihood including the GMRF, we must further include terms corresponding to the differences *ψ*_*a*(*i*)_ − *ψ*_*i*_. We do this by constructing matrices with block partitions for the coalescent likelihoods and GMRF likelihoods:

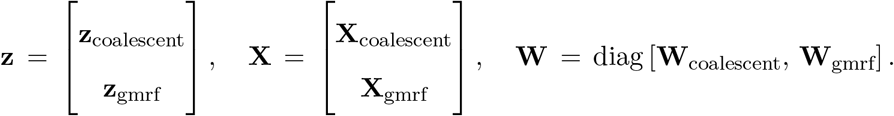

Here **z**_gmrf_ is zero for all rows reflecting smoothness assumption of the GMRF, **X**_coalescent_ has one 1 per row in the column of branch *i*, and **X**_gmrf_ has +1 in column *a*(*u*) and −1 in *u*. The weight matrices are diagonal with elements:

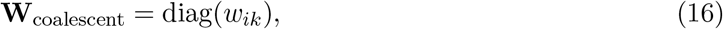

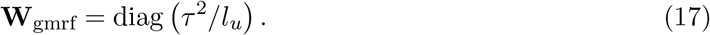

Solving the following weighted least squares problem then provides the estimates of *ψ*:

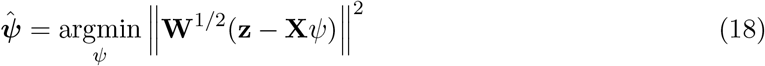

This yields matrices which are very large, but also very sparse and therefore can be inverted easily; thus solving these equations is much faster than maximising the original likelihood.

### 2.4 Optimising precision parameters

In general, models for continuous evolution of traits on a tree require the specification of parameters which control how correlation of the trait varies with evolutionary distance. The preceding sections were premised on the precision parameter *τ* being given. Specification of precision parameters is often carried out a priori or on an ad-hoc basis; choice of such parameters is a problem shared with many non-parametric statistical models, including spline regression and unsupervised clustering. We therefore develop solutions for optimising *τ* that are similar to approaches that have been used in a time-series regression context, and which leverage the fact that our objective function may be interpreted as a metric of predictive accuracy. For example, the likelihood (Eq. 7) is akin to the product of probabilities of new samples coalescing with a given branch; this likelihood is maximised when it is known exactly with which branch a new sample will coalesce. We may therefore select *τ* using time series cross validation techniques similar to those used in statistical forecasting methods [20]. This works by optimising the predictive accuracy by partitioning data into retrospective cohorts, fitting the coalescent odds model, and evaluating the objective function with new data from subsequent cohorts.

We implemented methods for calibrating *τ* based on the squared loss in Eq. 18 (Algorithm 1). Cohorts *j* with upper time-limit *c*_*j*_ are defined by collecting all nodes *i* with *t*_*i*_ < *c*_*j*_. The number of descendants in the following period *y*_*ij*_ are tabulated for each node and cohort. The particular times *c*_*j*_ of each cohort are determined by selecting quantiles *p*_*j*_ with a stratified random sample between user-defined limits. By default, we select cohorts in the latter quartile of *t*_*i*_ so that *τ* is well calibrated to recent evolution. The weighted least square (WLS) estimator is fitted to data in each cohort C_j_ and then the quadratic loss from Eq. 18 is computed for data in *C*_*j*+1_ \ *C*_*j*_. A grid-search followed by univariate optimisation (golden-section search) is used to minimise the loss with respect to *τ*.

#### Algorithm 1

Optimizing smoothing parameter *τ* given 1) a phylogeny *T* and 2) a method (e.g. the WLS estimator) for estimating coalescent odds *f*: *T* → *ψ*.

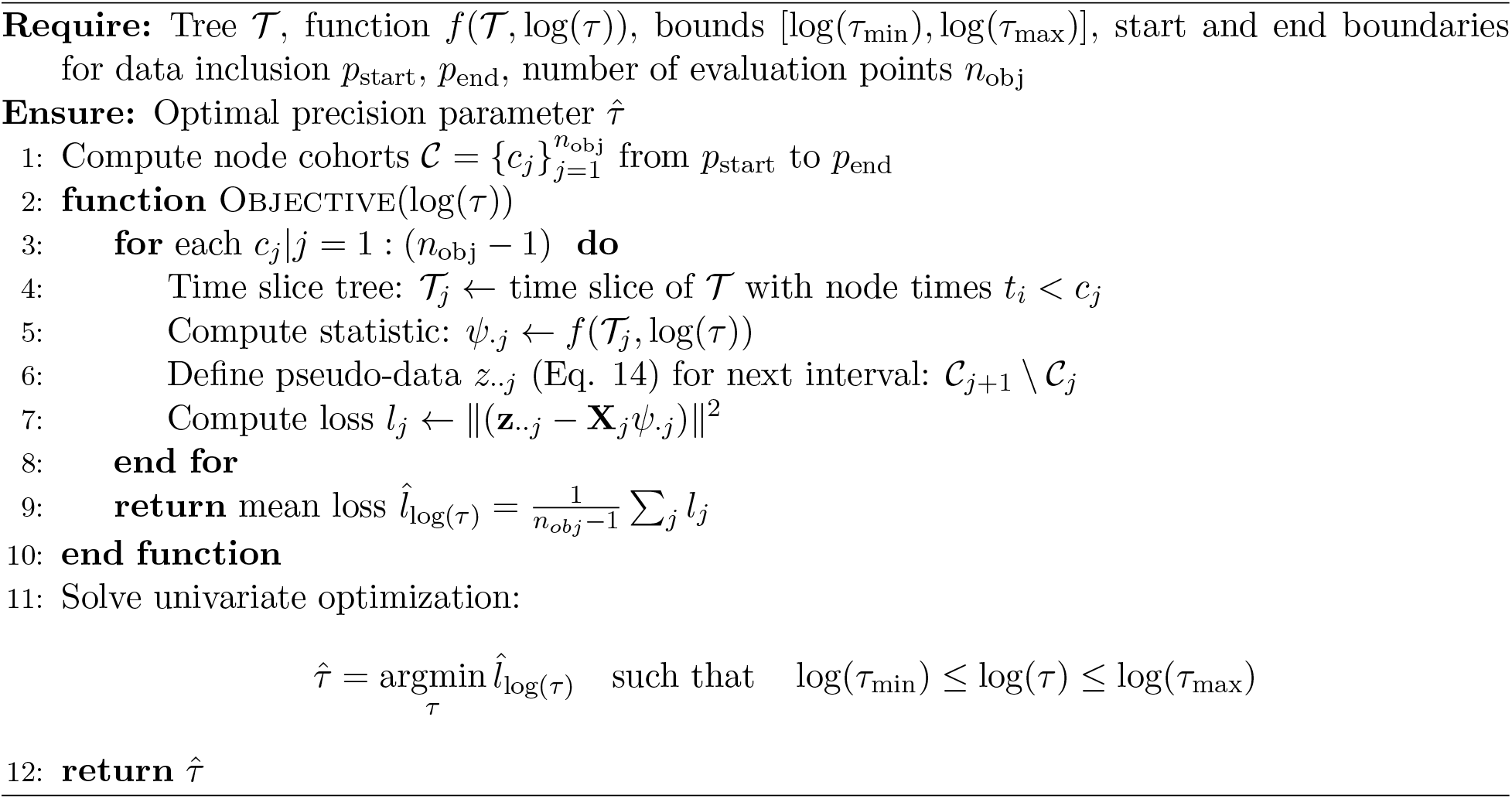

### 2.5 Detecting and adjusting for biased sampling

Whereas non-uniform sampling can skew coalescent odds towards populations with greater sample inclusion probabilities, we can easily adjust the estimated coalescent odds by modifying weights in the weighted least squares estimator (Eq. 18).

Suppose we have auxiliary data *Z*_*i*_ for each sample *i* that is associated with sample inclusions probabilities *π*_*i*_. Further suppose that these variables are potential confounders, and do not have any mechanistic impact on lineage fitness or growth, influencing only sampling patterns. For example, *Z* may include the geographic region for each sample unit, demographic information, or host-species. First, consider the case where *π*_*i*_ is a deterministic function of *Z* and assume *Z* is known. This may be the case, if for example different public health authorities during a pathogen outbreak have known sampling proportions, and we know the health authority associated with each sample.

In this case, we adjust the weights to take the new form with inverse probability weights:

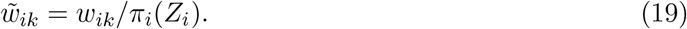

The revised estimator is obtained by substituting 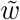 for *w* in Eq. 18.

Note that this adjusted weight only works if we know *π* for each node *i*, which in the best case is only true for tips of the tree. For internal nodes (ancestral to the sample), sample inclusion probabilities are not observed or known a priori, and we must estimate these based on available information at the tips of the tree. Ancestral state estimation of continuous variables (the *π*_*i*_) can be accomplished in numerous ways [21, 22]. We have implemented a simple, common and fast Brownian motion model of the ancestral *π* which is fitted using a least squares approach [23, 24].

For genomic epidemiology applications, it is often the case that the functional relationship between *π* and *Z* is not known, but there are nevertheless auxiliary variables which are hypothesised to influence sample inclusion. For example, we may know the time and location of each sample, but not have detailed information about how sampling rates varied over time and space. In this situation, we may be concerned about erroneous detection of lineages with high coalescent odds, and to reduce the false detection rate, we can calibrate *π* to achieve the most conservative (smallest in absolute value) *ψ* for lineages of interest.

Specifically, suppose *Z* is a binary variable taking values 0 and 1, and *π* has two values corresponding to these levels. And let *π* = *e*^−*βZ*^, so that the scalar *β* calibrates the degree of preferential sampling on *Z* = 1. We test the null hypothesis that *ψ* has equal mean across levels of *Z, H*_0_: E[*ψ*_*i*_(*β*)|*Z*_*i*_ = 1] = E[*ψ*_*i*_(*β*)|*Z*_*i*_ = 0], and quantify the strength of the association in terms of the t-statistic *T* (*β*), expressing the dependence of the association on sample weights. If the null is rejected, it implies that *Z* is influencing the results. We therefore calibrate *π* by selecting the minimal weight that removes the association of *ψ* with *Z*:

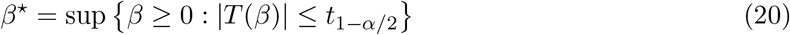

This procedure mitigates preferential sampling by attributing any association between *Z* and *ψ* to oversampling, and also proposes the degree of sample bias required to produce such an association.

### 2.6 Detecting phylogenetic clusters

A tree-cutting algorithm [25] combined with estimated coalescent odds can provide a convenient calculation of phylogenetic clusters – sets of samples that are connected by phylogenetic branches that do not have large intra-cluster differences in coalescent odds. This can be useful for identifying clades of closely related samples with similar growth rates and coalescent patterns.

The tree-cutting algorithm is based on the difference in coalescent odds between each node and its parent:

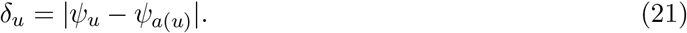

A threshold difference *ω* is used to cut the tree at the subset of nodes (both internal nodes and tips):

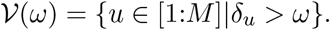

Let *D*(*u*) denote the set of nodes (including tips) descended from node *u*. The cluster corresponding to a node *u* in *V* is then the set of samples descended from *u* and not descended from any other node in *V* that is descended from *u*:

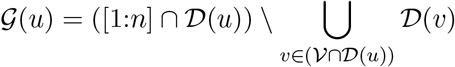

In practice, these clusters are easily computed given *V* and by doing a single post-order traversal of the tree, keeping track of all *D*(*u*) meeting the criterion above.

The composition of clusters will depend on the threshold change in coalescent odds *ω*, and the choice of threshold is ultimately subjective and could be tailored to various tasks. For example, a small threshold *ω* may be appropriate for the task of developing a very sensitive early warning signal for infectious disease surveillance (at the cost of lower specificity), where new clusters may represent an outbreak in its early stages. Nevertheless, there are some heuristics for choosing useful clustering thresholds in the absence of other data, and we have implemented an approach based on maximising the Calinski-Harabasz (CH) index [26]. Let *κ*(*ω*) denote the number of distinct clusters given threshold *ω*. We define the CH index in terms of the coalescent odds and given a threshold *ω* as follows:

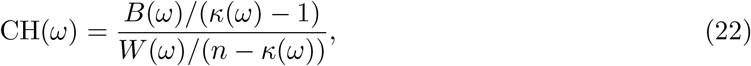

where *B*(·) and *W* (·) are respectively the between-cluster and within-cluster dispersions of the coalescent odds:

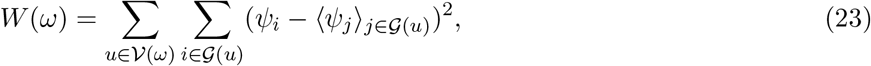

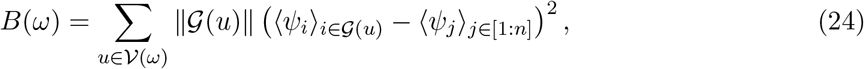

where ⟨·⟩_*X*_ represents the arithmetic mean over set *X*.

### 2.7 Implementation and simulations

Software to estimate coalescent odds is implemented as an open-source R package cod [27].

We developed simulations to evaluate the utility for coalescent odds estimated by least squares to identify lineages with a fitness advantage. We simulated two populations, an ancestral variant with population size *N*(*t*), and a variant with size *V*(*t*) introduced at time *t*_*v*_. Both types grow logistically with shared carrying capacity *K*, but the variant has a higher reproduction rate by a factor of 1 + *s*:

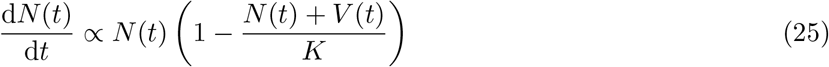

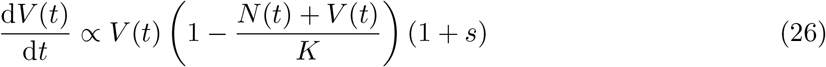

We simulated genealogies based on this model using a structured coalescent model implemented in the *Coalescent*.*jl* library [28]. Sampling was heterochronous over time (*n* = 1000) over 104 generations (equivalent to 2 years with *T*_*g*_ = 1 week), and constant sampling through time to mimic real sentinel surveillance data. The sample was additionally enriched with 100 random samples at the final time point *t*_final_. The ability to detect a deviation in coalescent odds amongst variant lineages depends primarily on the selection coefficient *s* and the relative prevalence of the variant at the time of the final sample *V*(*t*_final_)/(*N*(*t*_final_) + *V* (*t*_final_)). These two parameters were varied over a large range and coalescent odds were compared between ancestral and variant samples using a Wilcoxon rank sum test.

We additionally carried out an experiment with a variant with size *U*(*t*) that has no fitness advantage (*s* = 0) but which is sampled at a greater rate in order to evaluate the ability of this method to both detect unequal sampling and to automatically adjust for it. The variant moreover has a lower carrying capacity *K*_*u*_ < *K*_*a*_, so it will always be a minority in the population, but not necessarily in the sample:

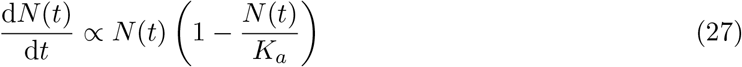

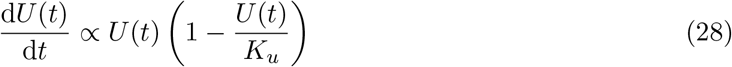

In all simulation experiments, the smoothing parameter *τ* was optimised as described in Section 2.4.

## 3 Results

### 3.1 Identification of high-fitness variants and adjusting for biased sampling

Simulations confirmed the expected dependence on both the size of the selective advantage and on the prevalence of the variant on the utility of coalescent odds for detecting lineages with a fitness advantage. Figure 1 illustrates the result of 2601 simulations over a range of values for the selection coefficient and variant prevalence parameters. It is possible to detect a fit variant even when the fitness effect is quite small (*s* = 0.01) or when the variant prevalence is quite small (5%), however detection under these conditions requires a large variant sample size and a large effect size, respectively. For context, during the COVID-19 pandemic, all major variants of concern had selection coefficients far in excess of the values simulated here, and sample density was quite high [29], suggesting that this method would be very sensitive for detecting similar variants even when they are at very low prevalence.

**Figure 1:**
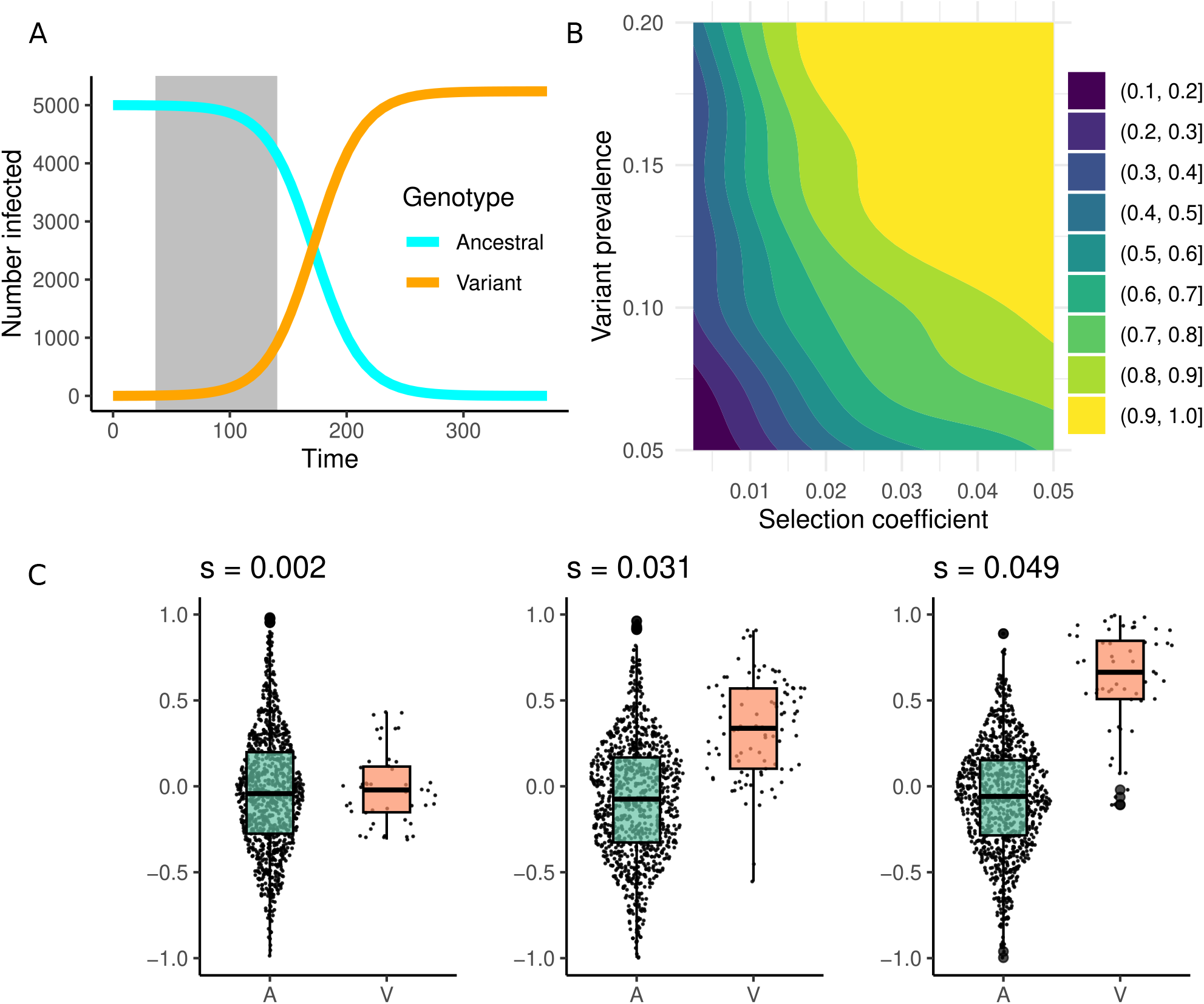
Estimation of coalescent odds in simulated populations with the emergence of a high-fitness variant. A. Population size of the ancestral and variant types over time. The shaded region indicates the period when sampling of both types takes place, ranging from 5% to 20% prevalence of the variant type. B. The probability that a significant difference in coalescent odds would be detected (1-tail Wilcox test *p* < 0.05) as a function of the selection coefficient in the variant type and the prevalence of the variant at the time of the final sample. Results are based on 2500 simulations. C. A comparison of coalescent odds amongst terminal branches of ancestral and variant types in three simulations with different selection coefficients. In all cases, the final sample is collected when the variant has 20% prevalence.

Regarding the sensitivity of coalescent odds to biased sampling, we find that it is relatively robust in terms of magnitude of effects detected, although significant differences are seen when the variant is sampled at more than twice the rate of the ancestral type. Figure S1 shows the distribution of coalescent odds amongst sampled lineages as the sampling weight is varied from two to ten. The magnitude of the differences in coalescent odds remains small (< 0.05) across scenarios. Adjusting the estimated coalescent odds using the algorithms described in Section 2.5 restores performance. Figure S2 shows p-values from a Wilcoxon rank sum test both with and without using automatically-tuned sample weights. These p-values remain well above common significance thresholds even when sampling the variant 10-fold.

### 3.2 Comparison of alternative estimators

Three objective functions were developed in the preceding sections to estimate coalescent odds: the likelihood (Eq. 8), the binomial likelihood approximation (Eq. 10), and the weighted least squares approximation (Eq. 18). Here we compare the coalescent odds estimated by each approach using a subset of simulations from Section 3.1. So that the comparison only features simulations where there is clear and strong contrast in coalescent odds between ancestral and variant types, we study the set of simulations with a selection coefficient *s* = 0.2 and prevalence of the variant > 15%. These are shown in Figure 2. In order to make the experiment computationally tractable for all methods, we further downsample to *n* = 150 with preferential inclusion of the more recent samples.

**Figure 2:**
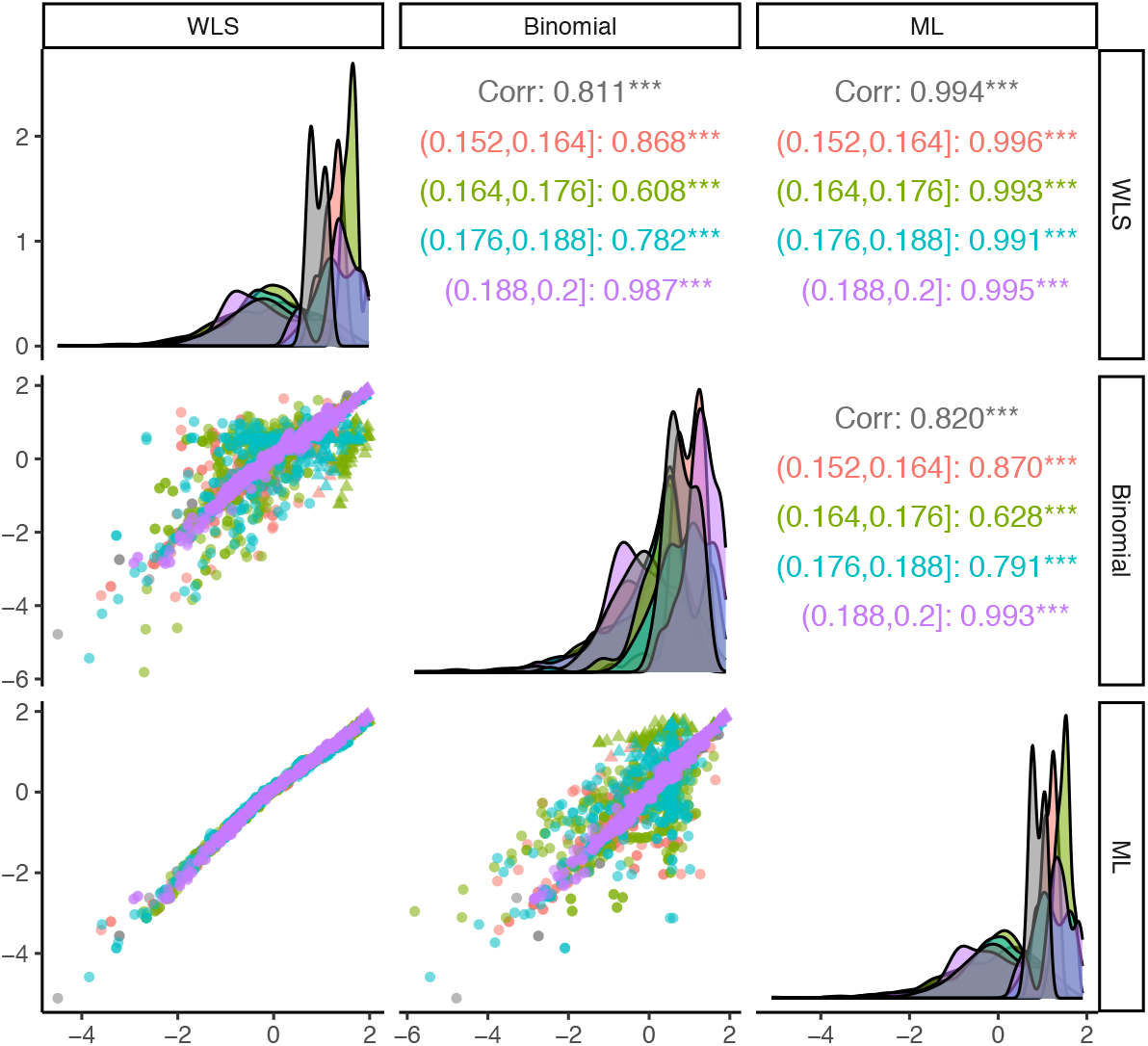
Comparison of coalescent odds estimated by least squares, a binomial GAM, and maximum likelihood. The lower off-diagonal panels show scatter plots of all possible combinations of estimators. The diagonal panels show kernel-density estimates of the coalescent odds for each estimation method. The upper off-diagonal panels show the Pearson correlation coefficients for each combination of estimation methods. The comparisons are further stratified by the prevalence of the variant type at the final time of sampling (colours). Correlations for all data combined are shown in black.

Overall, we found extremely close correspondence between the maximum likelihood estimator and the weighted least squares approximation supporting the use of the 1-step IRLS approximation. The Pearson correlation between these estimators exceeds 99% across all replicates and scenarios. The computation time is drastically shortened by the use of this approximation (more than 10 thousand fold in this simulation set).

The greatest contrast between methods is seen between the true likelihood and the binomial likelihood approximation. The latter shows much higher dispersion in estimates when prevalence of the variant type is low. The Pearson correlation in this case falls as low as *ρ* = 63%. We also find negligible computational benefit to the use of this approximation (40% speedup), although the implementation in the `mgcv` package would easily support further extensions using covariate data and more complex correlation structures.

### 3.3 Application to antimicrobial resistant lineages

We analysed a dataset of 1,102 genomes of *Neisseria gonorrhoeae* from the United States [30]. A time-scaled phylogeny was constructed by first using phyml [31], then correcting for recombination using ClonalFrameML [32] and finally dating the nodes using treedater [33]. The coalescent odds (*ψ*) were estimated along branches using the weighted least squares approach, which took about one minute on a standard laptop computer. Smoothing parameters were optimised as described in Section 2.4. We assessed potential sampling bias across clinics using the methods in Section 2.5; the association between the coalescent odds and key location of sampling persisted even under extreme weights, indicating the observed signal is unlikely to be an artifact of over-sampling. We then modeled the coalescent odds against resistance status and time across six antibiotic classes and projected their values at the end of sampling. As shown in Figure 3, only azithromycin (AZI) resistance is associated with elevated coalescent odds, whereas other antibiotic classes show no comparable positive effect.

**Figure 3:**
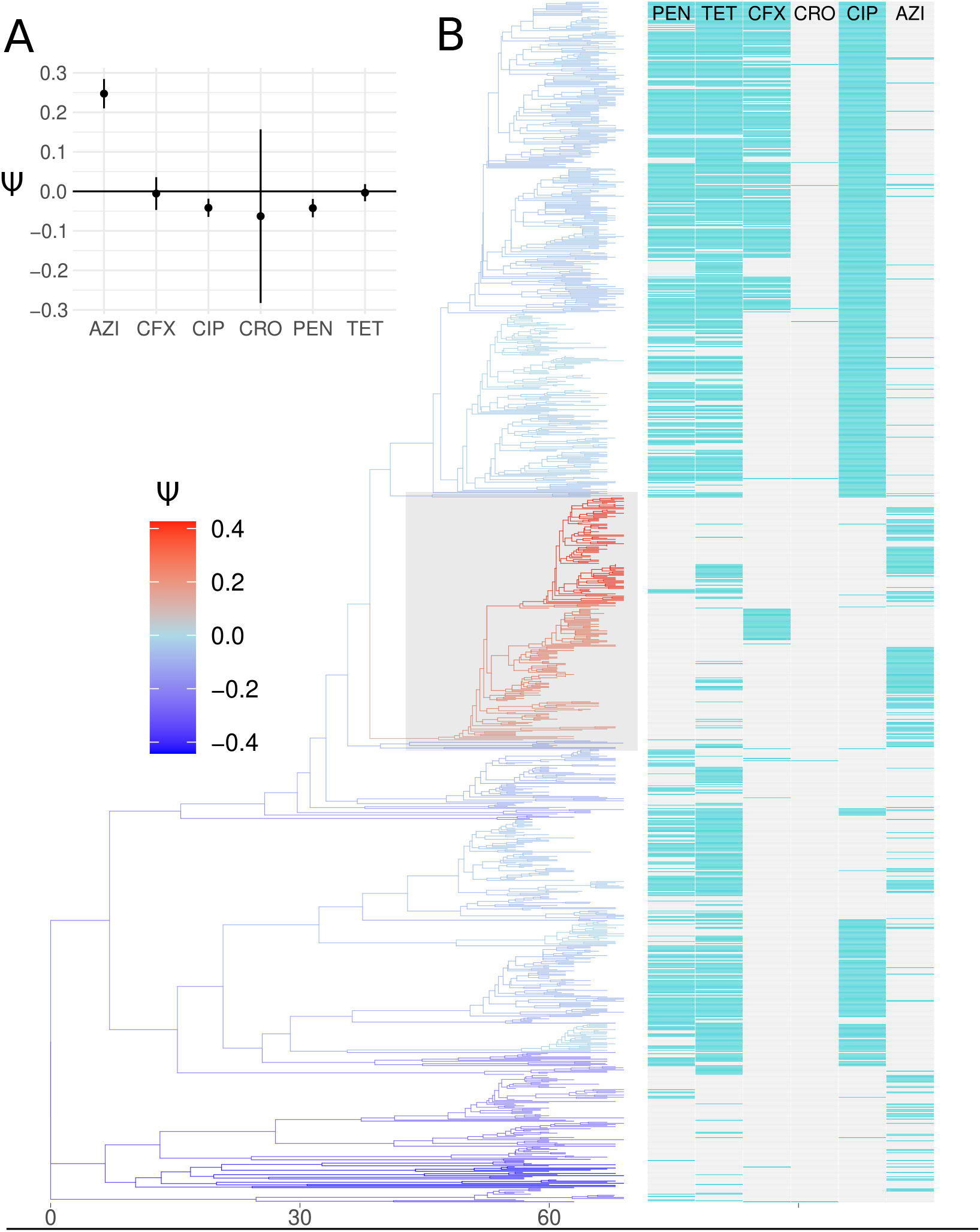
Coalescent odds estimated for a phylogeny reconstructed from 1,102 *Neisseria gonorrhoeae* whole genomes collected in the United States between 2000 and 2013. The estimates are stratified by antibiotic resistance phenotypes (presence/absence) for: penicillin (PEN), tetracycline (TET), cefixime (CFX), ceftriaxone (CRO), ciproflaxicin (CIP), and azithromycin (AZI), and resistance phenotype is shown for each sampled lineage. A. The mean and 95% confidence interval for the estimated coalescent odds of sampled lineages in 2013 stratified by antibiotic resistance phenotype. B. Time-scaled phylogeny coloured by estimated coalescent odds. The shaded clade was detected by the cluster-detection method.

To summarise growth at the clade level, we defined *ψ*-based phylogenetic clusters by thresholding lineage-wise changes in *ψ* as described in Section 2.6, selecting the threshold via the Calinski-Harabasz index [26]. This revealed a prominent cluster with high coalescent odds that closely corresponds to AZI-resistant isolates (Figure 3B). Thus, both continuous estimates of the coalescent odds and the definition of clusters based on their values converge on the same signal of a clade that is both growing and featuring elevated levels of AZI resistance.

Notably, the most recent time point in this dataset showed < 1% AZI resistance in 2013 and time series data to this point did not show an obvious signal of growth in AZI resistance. However, by 2017 it had grown to roughly 5% [34]. This rapid expansion is consistent with the elevated *ψ* inferred for AZI, supporting the use of the coalescent odds as an early indicator of lineages likely to grow.

### 3.4 Comparison to local branching statistics and SARS-CoV-2 reproduction numbers

To assess the utility of coalescent odds as a lineage detection tool for emerging variants, we reanalysed data from a recently published method, phylowave [35]. This method was shown to reliably detect major lineage dynamics and selective sweeps for SARS-CoV-2, seasonal H3N2 Influenza virus, and *B. pertussis*. The phylowave method employs a modified and well-calibrated version of the local branching index [3], as an initial filter for detecting significant clades. We therefore computed coalescent odds and the phylowave index for the three time-scaled trees previously analysed in the phylowave study [35] and directly compared their values at tips in each tree.

Computation of the phylowave index requires manual calibration of four hyperparameters: the mutation rate, genome length, a kernel timescale, and a time window over which the statistic is computed. Whilst the first two parameters are evident from the data, the latter two require expert judgement and can shape the output. We re-used values of these parameters from the original study. When computing coalescent odds, we used the weighted least squares approximation with default settings, which automatically selects the smoothing parameter using Algorithm 1. Manual selection of hyperparameters was not required.

Figure 4 shows the estimated coalescent odds and phylowave index for the SARS-CoV-2 dataset (*n* = 3, 129) which spans the pandemic period. Supporting figure S3 shows the comparison with the other datasets. We observe that the statistics are capturing different aspects of lineage replacement. Each major lineage manifests a discrete change in coalescent odds, which correlates with the major change in fitness seen with each selective sweep. The phylowave index in contrast has a boom-bust pattern that correlates with the initial growth of each major lineage. The correlation coefficient between the statistics is actually quite low (Pearson r=7%), but this is attributable to the strongly time-dependent nature of the phylowave index. Similar patterns were seen for H3N2 and *B. pertussis*.

**Figure 4:**
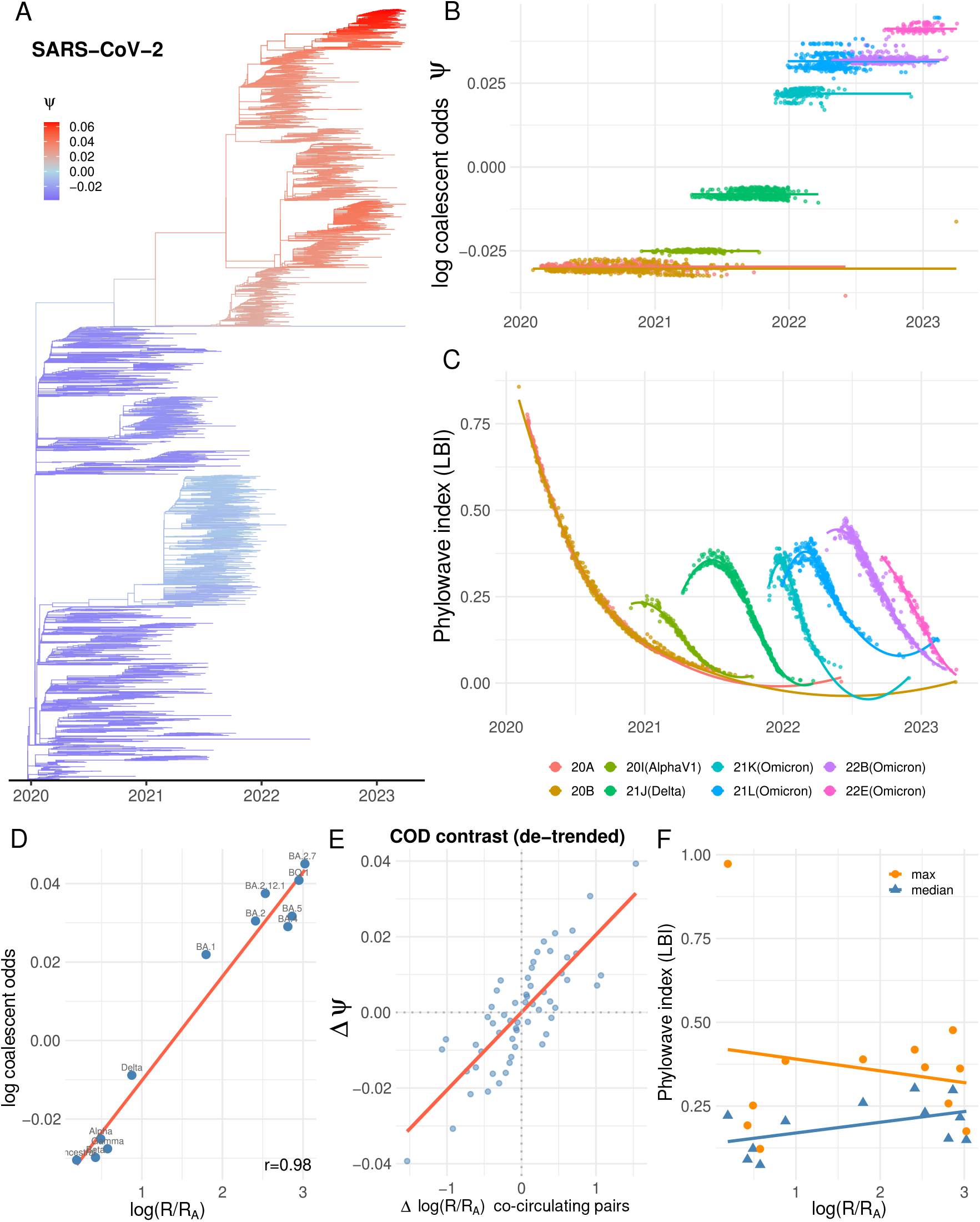
Coalescent odds and the phylowave index for a SARS-CoV-2 phylogeny reconstructed from global samples 2020-2023 (*n* = 3129). A: Time-scaled tree from [35] showing log coalescent odds. B and C: Coalescent odds and phylowave index for the main Nextstrain clades with a smooth regression line (colour). D, E and F: Coalescent odds and phylowave index versus reproduction numbers of each clade estimated in [36, 37]. E shows the difference in log coalescent odds versus the difference in reproduction numbers considering all pairs which co-circulate.

To elucidate how these statistics are associated with the fitness of each variant, and the magnitude of selection driving each selective sweep, we further examined the relationship between coalescent odds, the phylowave index, and independent estimates of the reproduction number [36] estimated using pyro-cov [37] (data up to September 2022). This allowed us to compare coalescent odds with a measure of fitness of each variant *i*, the ratio of the reproduction number of that variant *R*_*i*_ with the reproduction number of the ancestral type (Wuhan) *R*_*A*_. We then use *F*_*i*_ = log(*R*_*i*_*/R*_*A*_) as our measure of replicative fitness for comparison with both statistics. The coalescent odds were computed under default settings and automatic hyperparameter tuning. No attempt was made to adjust for unequal sample intensity amongst different variants. Using the regression mean of *ψ* for each variant, we find that *ψ* and log(*R*_*i*_*/R*_*A*_) are strongly associated (Pearson *r* = 98%, Figure 4D). Since some correlation between *ψ* and *F*_*i*_ is inevitable given the sequential and strictly increasing nature of the series, we applied a more stringent test, comparing the difference in coalescent odds between variants to the difference fitness (Figure 4E). Note that each point corresponds to *F*_*i*_ − *F*_*j*_ = log(*R*_*i*_*/R*_*j*_) for cocirculating clades *i* and *j*. These data included all pairs of Nextstrain clades which cocirculated in time after removing outliers (0.5% tails of their respective sample time distributions). This comparison retains a very strong linear relationships (Pearson *r* = 0.81, permutation test *p* < 10^−3^) and demonstrates that the difference in *ψ*_*i*_ can be interpreted as a proxy for the relative fitness in individual selective sweeps.

Elucidating the relationship between the LBI and fitness is more challenging because this statistic is highly changeable; we therefore examined both the maximum phylowave index, which would plausibly be positively associated with the initial rate of lineage growth, as well as the median Phylowave index. These statistics did not correlate with *F*_i_ (maximum index *p* = 0.57 and median index *p* = 0.11), since they do not have an increasing trend (Figure 4E) as *ψ* does. We therefore looked for an association between the phylowave index and *F*_*i*_ − *F*_*j*_ for the subset of *i* and *j* corresponding to selective sweeps: Ancestral→ Alpha → Delta, Delta → BA.1, BA.1 → BA.2, BA.2 → BA.5, and BA.5 → BQ.1. This showed larger correlation (max index *r* = 0.305 and median index *r* = 0.47 as compared to *r* = 0.738 for Δ*ψ*), but possibly because there were only six contrasts, this fell short of significance for all statistics including Δ*ψ*.

## 4 Discussion

We have presented a novel coalescent process where the propensity to coalesce varies continuously as a heritable trait (Section 2.2), and we developed several methods to fit this model to phylogenetic data. We further developed a weighted least squares estimator which greatly reduces computational effort on large trees (Section 2.3); and, we present new algorithms for calibrating hyperparameters using a time-ordered cross-validation to select parameters which maximise predictability of evolution (Section 2.4). It is relatively fast to implement in combination with the weighted least square approximation. The clustering methods presented in Section 2.6 provide a convenient way to leverage coalescent odds to automatically identify important clades for subsequent analysis.

When comparing coalescent odds with branching indices (Section 3.4), we observed that they measure different but related dynamics. Branching indices show periods where lineages grow quickly, but are computed locally in tree space, so that differences with co-circulating lineages are not quantified if they are not closely related. In contrast, coalescent odds are computed relative to contemporaneous lineages even if they are distant in the tree, so that durable changes in pathogen fitness are estimated. Measuring the strength of selection requires calibration relative to contemporaneous lineages, not just close lineages, and coalescent odds have utility as a tree-based proxy for selection. Either type of statistic could be used for monitoring and detecting emerging variants, and future work should explore the relative sensitivity and power of these methods to produce such signals, although our current results suggest that coalescent odds are more strongly associated with lineage fitness if estimated retrospectively (Section 3.4). Branching indices have an advantage in computational speed, however they are dependent on expert judgement of several hyperparameters. This can be difficult for emerging pathogens where there is limited prior knowledge. A potential advantage of coalescent odds in this context is that the statistic is calibrated with a single hyperparameter and we have proposed efficient algorithms for automatically calibrating it.

In simulations, we observe that estimation of coalescent odds is a highly sensitive method for detecting lineages that are expanding due to positive selection. The weighted least square estimator is moreover scalable to relatively large datasets with more than a thousand samples. These findings are born out by our re-analysis of 1,102 *Neisseria gonorrhoeae* genomes. This illustrates the potential utility of these methods as an epidemiological surveillance tool, especially when combined with important phenotypic or clinical variables. For example, this method indicates a rapidly growing clade with elevated azithromycin resistance, and it is retrospectively clear that this signal could have been detected in the phylogenetic analysis before azithromycin resistance grew to noticeable levels [34].

Non-uniform sampling can lead to over-representation of closely related lineages and consequently higher coalescent odds associated with oversampled populations. This is a common problem in genomic epidemiology [38, 39], where convenience sampling is common, or sampling is undertaken in response to an outbreak. Furthermore, most epidemics have a geographic range that spans multiple public health authorities, and each will have different capacities for sampling and sequencing. Currently, there are not any lineage growth statistics or phylogenetic clustering methods that can account for sampling variability.

Regarding the implications of biased sampling for false-discovery of lineages with high fitness, this method provides a substantial improvement over current methods. We find, as expected, that if sampling is skewed towards a lineage it will have higher coalescent odds. But we also find that the estimation of coalescent odds is relatively inelastic to variation in sample weights, so that the magnitude of the bias is small relative to the effects stemming from real differences in fitness. If sample weights are known, these can be applied in a fashion that is similar to how they are used in regression analysis. If sample weights are not known, but if we have an auxiliary variable which we suspect is associated with sampling, we also propose a method to automatically tune sample weights to give the most conservative possible estimate of coalescent odds (Section 2.5).

Estimation of coalescent odds is currently limited to datasets where an accurate time-scaled phylogeny can be estimated. We have not examined the sensitivity of these estimators to phylogenetic error and molecular clock dating error. However, results are likely to be unreliable if the time-scaled phylogeny is grossly misestimated, as for any post-processing method [40]. Future work may explore relaxation of this constraint; similar models may be developed in cases where genetic data do now allow robust phylogenetic inference (e.g. due to slow molecular evolution) but where there is very complete sampling over time for a long duration and genetic clustering is still possible.

There is very wide scope to improve on this category of models and inference methods. We have demonstrated how metadata can be used to weight sample units, but it would be a relatively simple extension to include covariate data to improve estimation of coalescent odds. For example, any variable that may be associated with the relative fitness of a taxon could be useful, and inclusion of such variables in the regression framework would provide a way to quickly combine virological data [41], computational estimates of fitness [42], and population genetic estimates of fitness [43] into a single framework. Although coalescent odds do not provide a direct estimate of fitness, future theoretical developments may illustrate how these concepts are quantitatively related, as has been recently carried out with the LBI and birth-death models in special cases [44]. Nevertheless, coalescent odds allow rapid detection of variants with a fitness difference, and other methods can then be applied [29]. For example, in the case of antibiotic resistant lineages, it could involve modelling explicitly the fitness cost and benefit of resistance to explain differences in phylodynamic trajectories [45].

There is also scope to relax assumptions underlying the current model for coalescent propensity, which is premised on a simple GMRF model, and to propose generative models to co-simulate trees and coalescent rate variation [2]. For large trees representing deep evolutionary time, the assumption of continuous and uniform jumps in coalescent propensity may be invalid. Future work should explore more complex hierarchical models, or perhaps leverage the current clustering method to partition and iteratively analyse clades of the tree which are likely evolving under different regimes.

## Acknowledgements

The authors thank Luca Ferretti, Ian Roberts, Megan Saathoff, and Chris Wymant for helpful feedback. EV acknowledges support from the UK Health Security Agency CARAA 104683ED “Development of phylogenetic analysis tools to track HIV transmissions for public health surveillance purposes” and CARAA 5126118 “Provision of expert advice and development support for emerging infections genomics and metagenomics analysis”.

## Data Availability Statement

Code to reproduce simulation data and the SARS-CoV-2 analysis are available on Zenodo at https://doi.org/10.5281/zenodo.20938043. The *Neisseria gonorrhoeae* phylogeny and metadata are included as a vignette in the cod R package [27].

