## Supporting figures for "Quantifying the predictability of evolution by analysis of coalescent rate variation"

Erik Volz Xavier Didelot

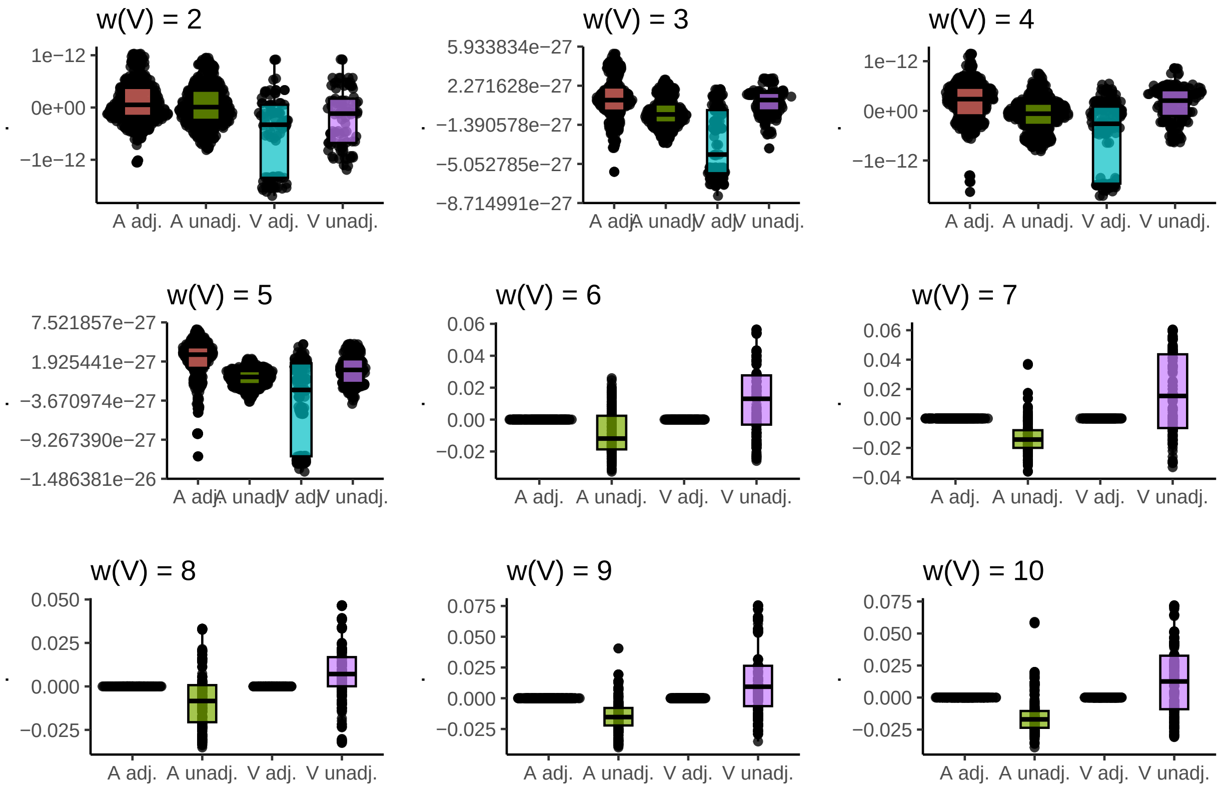


Figure S1: Comparison of coalescent odds between 1) Ancestral and variant types, and 2) with and without sample weight adjustments. Sample weights are varied from 2 to 10.

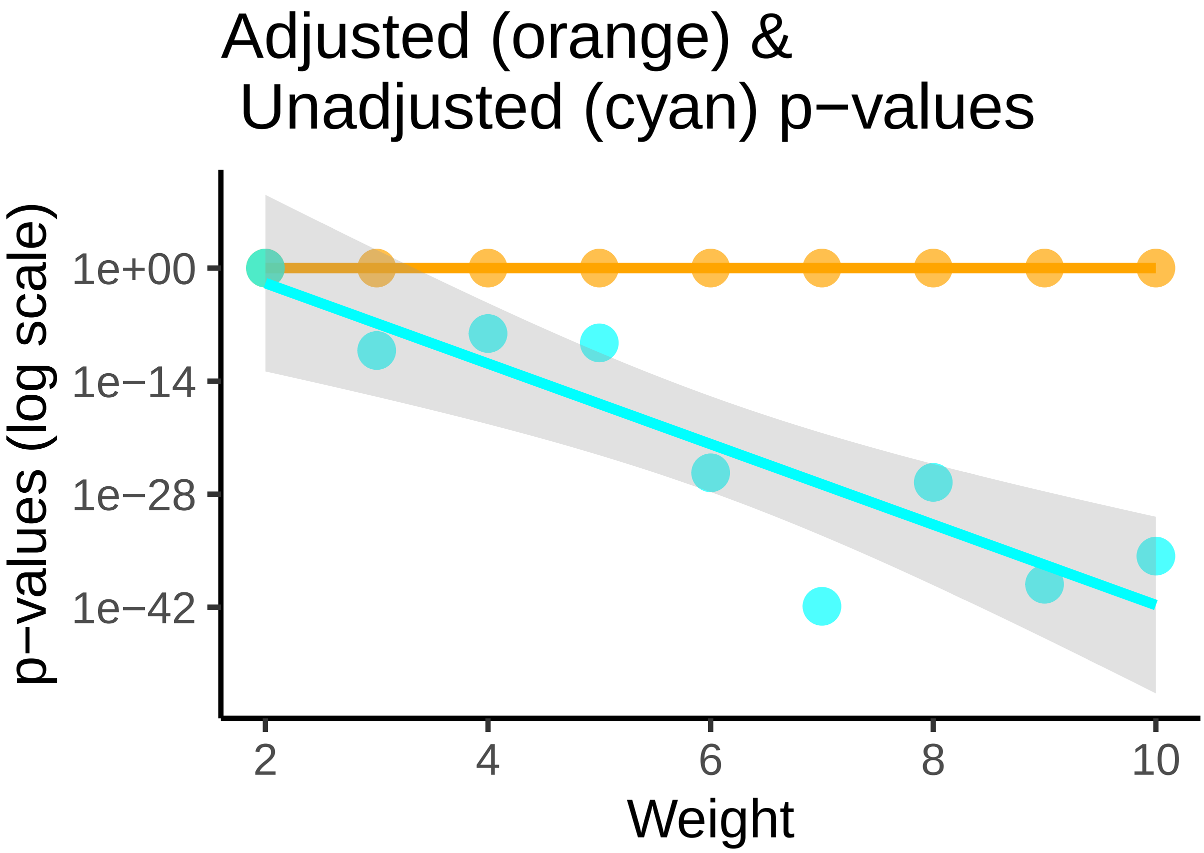


Figure S2: The p-values (log scale) of a Wilcoxon rank sum test comparing variant and ancestral coalescent odds (sampled lineages only), and comparing estimates with and without sample weighting with automatic tuning.

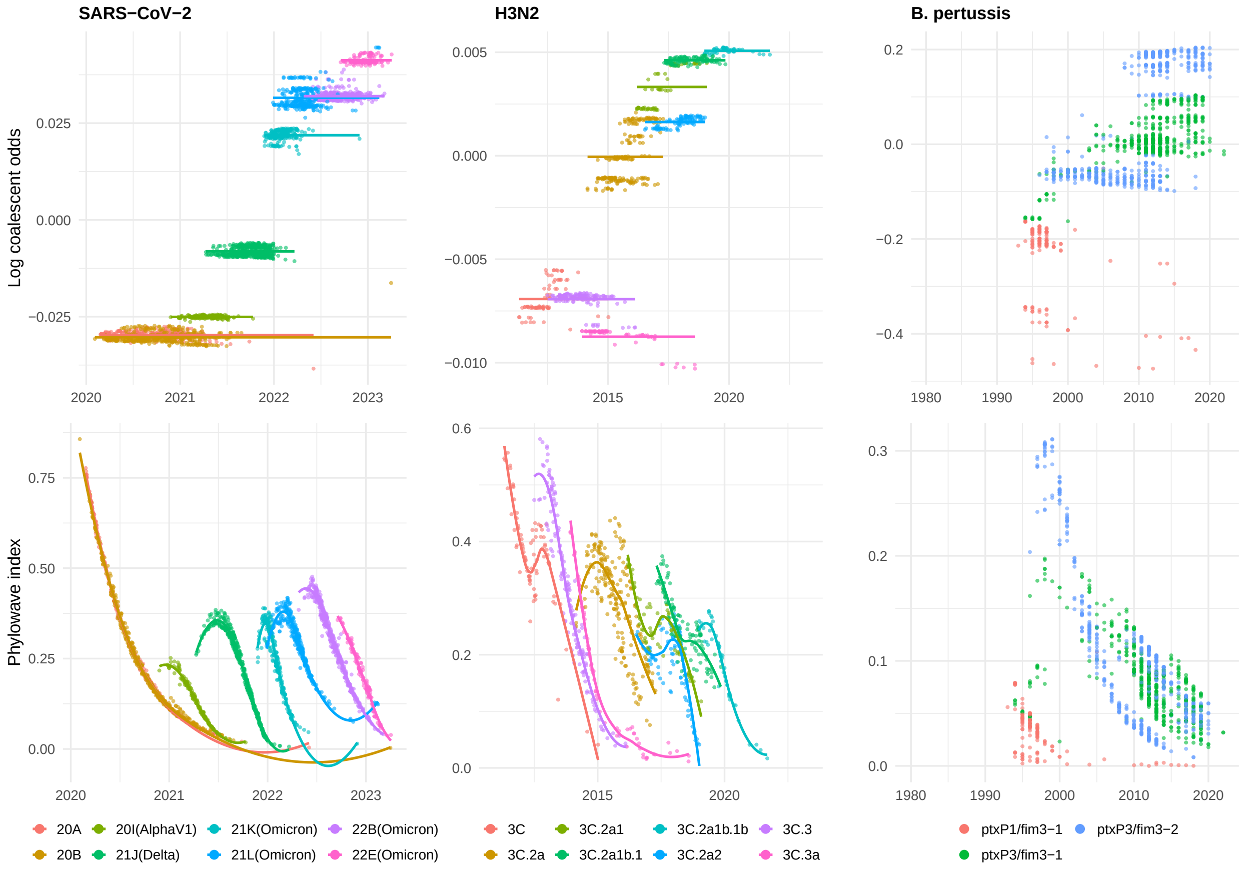


Figure S3: Comparison of coalescent odds (top, WLS approximation with default automatically tuned smoothing parameter) and the phylowave index (bottom, same parameters used in [1]). Only values from tips are shown.
